# Alternative splicing plasticity of the neurexin family of synaptic adhesion molecules in human sensory neurons

**DOI:** 10.64898/2026.09.02.748890

**Authors:** Judy J. Yoo, Bryan A. Copits

## Abstract

Plasticity within somatosensory circuits enables adaptations to environmental changes, however these can become maladaptive following injury and lead to chronic pain states. Presynaptic neurexins (Nrxns) are key organizers of synapse function that bind to a wide variety of postsynaptic proteins in a splice-isoform dependent manner to regulate synaptic connectivity, transmission and plasticity. Here we observed that *NRXN* alternative splicing at splice site 4 (SS4), an exon critical for synaptic function and long-term potentiation at synapses in the central nervous system, is plastic in human sensory neurons of the dorsal root ganglia (DRG). Taking a reverse translational approach, we used mouse models to understand what might regulate these splicing changes in DRG. We observed a shift to transcripts lacking this alternatively spliced exon for *Nrxn1* and *Nrxn3* in response to depolarization, but not sensitization with PGE_2_. Alternative splicing at SS4 was not altered by acute inflammatory pain but was reduced by central axotomy *in vivo*. We found that *Nrxn* alternative splicing patterns in DRG are dynamic during nervous system development, and vary between neurons of the DRG, spinal cord, and cortex in adult animals. Lastly, we observed that exon use at this site is conserved between species for Nrxn1α, but significantly different for all other isoforms. Thus, *Nrxn* alternative splicing at SS4 is plastic in human and mouse DRG neurons, which may influence somatosensory circuit wiring and synaptic function.

**Significance statement:** The neurexin family of adhesion molecules act as key organizers of synaptic connectivity and function. These molecules undergo extensive alternative slicing to generate thousands of isoforms that can influence their interactions with diverse families of postsynaptic adhesion molecules. We found that alternative splicing at one of these key sites is altered in human somatosensory obtained from organ donors with damaged spinal grey matter. We then took a reverse translational approach in mouse model organisms to understand if *Nrxn* alternative splicing is plastic in peripheral somatosensory neurons of the dorsal root ganglia.

## Introduction

Somatosensory information is encoded by ion channels and receptors in distinct neuron types of the dorsal root ganglia (DRG) where it is then transmitted to the central nervous system. While much is known about the identity of DRG cell types and their molecular transducers, it is less clear how these projections are wired to correct postsynaptic targets in the spinal cord and brain. Trans-synaptic interactions between diverse families of cell adhesion molecules are well poised to mediate both neuronal recognition during circuit assembly (de Wit and Ghosh, 2016; Sanes and Zipursky, 2020) and synaptic function (Biederer et al., 2017; Kim et al., 2021; Connor and Siddiqui, 2023) to enable adaptive responses to ensure survival. Owing to their critical roles in establishing appropriate neuronal connections, many of these synaptic adhesion molecule families have strong genetic linkages to human neurodevelopmental disorders (Chowdhury et al., 2021; Exposito-Alonso and Rico, 2022). Somatosensory alterations are highly prevalent in autism spectrum disorders (Foss-Feig et al., 2012; Balasco et al., 2020) suggesting that these may arise from miscommunication at the first sensory synapse (Orefice et al., 2016; Orefice, 2020).

The neurexin family of synaptic adhesion molecules consists of three genes (*Nrxn1- 3*) that utilize separate promoters to produce long α-Nrxn or shorter β-Nrxn transmembrane proteins (Ushkaryov et al., 1992; Ullrich et al., 1995; Gomez et al., 2021). The *Nrxn1* gene can also generate a very short y-isoform lacking the major extracellular domains found in α/β-Nrxns (Sterky et al., 2017). Over 12,000 unique variants can theoretically be generated from only three *neurexin* genes through the combinatorial use of different promotors and up to six alternative splicing sites (SS1-6) (Gomez et al., 2021). Single molecule long-read sequencing has confirmed over 5000 distinct *Nrxn* variants that are expressed in mouse brain (Schreiner et al., 2014; Treutlein et al., 2014) and alternative splicing patterns and exon usage for *NRXN1*a appear to be conserved in human neurons(Flaherty et al., 2019). This extensive diversity has been proposed to represent a molecular code for establishing neuronal connections and regulating synaptic function through *trans*-synaptic interactions with different postsynaptic cell adhesion molecules (Südhof, 2017).

Of these alternative splice sites, SS4 has been the most widely studied, owing to its strong influence on neurexin binding affinity to a diverse range of postsynaptic ligands including neuroligins (Ichtchenko et al., 1995; Scheiffele et al., 2000; Boucard et al., 2005), leucine-rich repeat transmembrane proteins (LRRTMs) (de Wit et al., 2009; Ko et al., 2009; Linhoff et al., 2009; Siddiqui et al., 2010), cerebellins (Matsuda et al., 2010; Uemura et al., 2010), dystroglycan (Sugita et al., 2001; Reissner et al., 2014; Trotter et al., 2023), and latrophilin adhesion GPCRs (Boucard et al., 2012). Functionally, SS4 splicing alters postsynaptic AMPA and NMDA receptor composition and plasticity in the hippocampus in a Nrxn isoform- and synapse-specific manner (Aoto et al., 2013; Traunmüller et al., 2016a; Dai et al., 2019; Trotter et al., 2023). These findings indicate that cell specific patterns of Nrxn isoform expression and alternative splicing may be central regulators of synaptic diversity throughout the nervous system. However, these studies have largely focused on well-established circuits in the brain, and the role of neurexins in somatosensory circuits are largely unknown. Moreover, direct comparisons of *Nrxn* alternative splicing between model organisms and human neurons are very limited, which hampers translating these findings toward a better understanding of human nervous system function.

Here, we set out to understand *NRXN* SS4 splicing profiles in human somatosensory neurons, hypothesizing that they may be altered in patients with prior pain history. While we did not observe any changes in *NRXN* splicing in DRG from these patients, we found a surprising shift towards SS4- transcripts in a subset of donors, suggesting that *NRXN* alternative splicing was plastic in human DRG. Taking a reverse translation approach, we turned to rodent models to understand what might drive this plasticity in human neurons.

## Results

### Neurexin alternative splicing in human somatosensory neurons

Both α- and β-*neurexin* (*Nrxn*) transcripts can undergo alternative splicing at site 4 (SS4) to include a 90 bp exon that influences synaptic connectivity and transmission (**Figure 1A**). We wanted to investigate alternative splicing of these presynaptic adhesion molecules at SS4 in human somatosensory neurons and test if this was altered in donors with reported pain histories. To assess alternative splicing at SS4, we designed a set of 5’ primers unique to the alpha and beta variants of all three human *NRXN* isoforms with common 3’ primers based on regions used previously in rodents (Kang et al., 2008; Iijima et al., 2011) to determine the relative proportions of SS4+ and SS4- transcripts (**Figure 1B**). We purified RNA from lumbar human dorsal root ganglia (hDRG) extracted from organ donors (Valtcheva et al., 2016) across a range of sexes, ages, and potential pain conditions, and performed RT-PCR followed by gel electrophoresis. We quantified the relative proportion of the larger PCR product containing this 90 bp exon at splice site 4 (SS4+) and divided this by the summed intensity reflecting total *NRXN* transcript expression to determine the relative proportion of exon splicing at SS4 (**Figure 1C-D**). We found that SS4 alternative splicing differed across NRXN isoforms in human sensory neurons, with greater incorporation of SS4 in the larger α-*NRXNs* compared to the shorter β-*NRXN* isoforms, which was most pronounced in *NRXN3)*. Overall, exon splicing at SS4 in human *NRXN* transcripts was similar across donors of different age and sex and was not altered in hDRG from donors with reported pain conditions (**Figure 1C-D**, **Table 1**).

**Figure 1:**
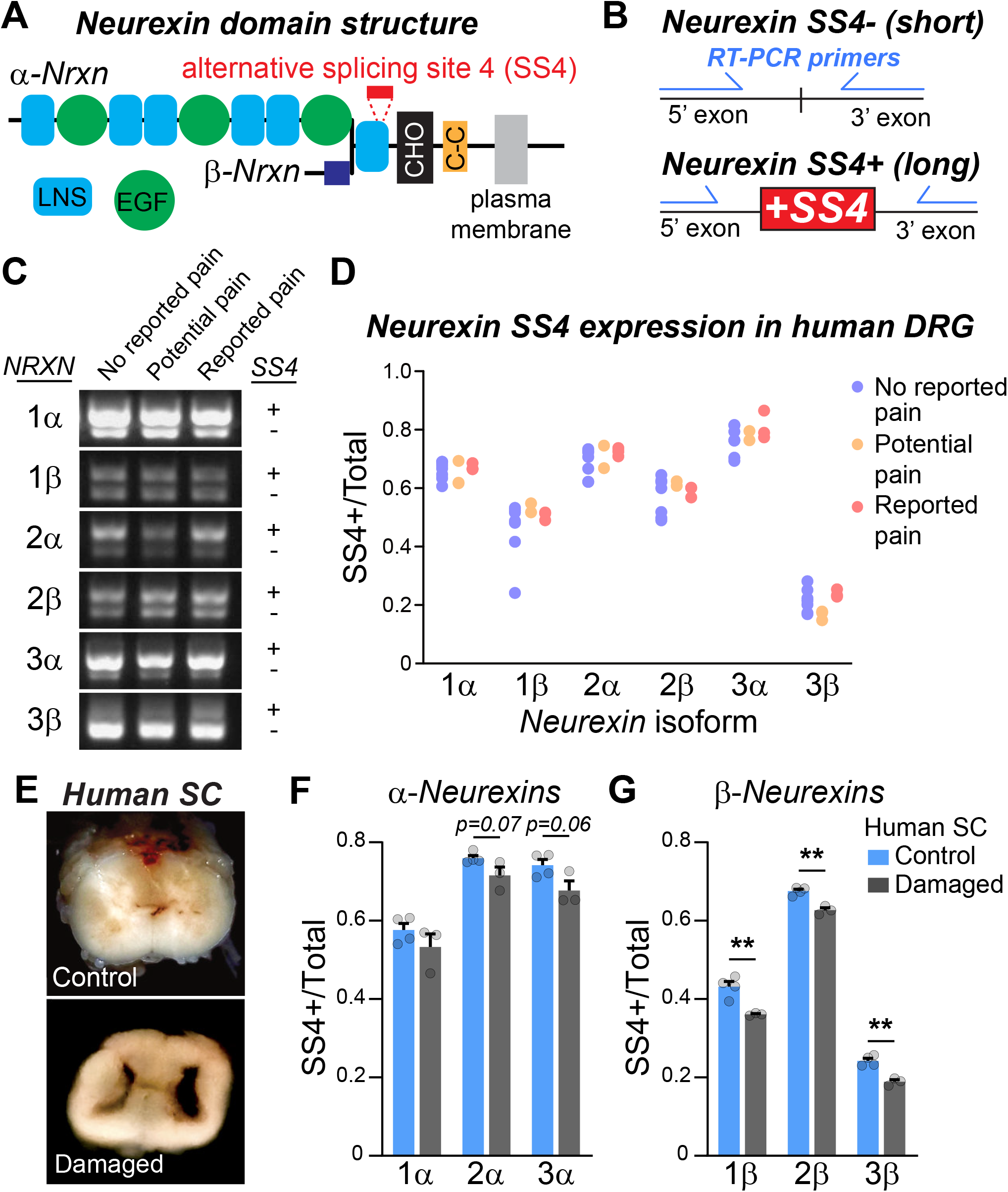
Neurexin alternative splicing in human somatosensory neurons. (A) Neurexin (Nrxn) protein domain cartoons for the larger α- and shorter β-Nrxn proteins. Alternative splicing site 4 (SS4) is a 90 bp insert in the shared LNS (laminin/neurexin/sex hormone binding globulin) domain on the extracellular surface of both variants. EGF (epidermal growth factor-like domain), CHO (O-linked glycosylation region), C-C (cysteine disulfide bond). (B) Diagram of RT-PCR strategy to detect Nrxn SS4 alternative splicing. (C) Images of ethidium bromide-stained gels from RT-PCR products of SS4+ and SS4- transcripts for both α- and β-*NRXN1-3* from human dorsal root ganglia (hDRG) obtained from three donors with differences in reported pain histories. (D) Summary graph of SS4+/Total *NRXN* splicing ratios for all neurexin isoforms in hDRG from donors with differences in reported pain histories. N=5 donors for no reported pain, n=2 with potential pain, and n=3 with reported pain history. (E) Images of transverse blocks of human spinal cord (SC) from two different donors with healthy vs necrotic gray matter. (F-G) Summary graphs of Nrxn SS4 alternative splicing for longer alpha (F) and shorter beta (G) isoforms from donors with necrotic gray matter vs. age- and sex-matched controls. N=3 donors with observed deterioration of the spinal gray matter and n=4 age- and sex-matched controls in which spinal gray matter was confirmed to be intact. **p<0.01 using unpaired t-tests.

**Table 1:** Donor demographics for human DRG used in this study. MVA: motor vehicle accident, ICH: intracerebral hemorrhage, CVA: cerebrovascular accident, SAH: subarachnoid hemorrhage, OD: overdose

| Pain history |  |  |  |  |  |  |
| --- | --- | --- | --- | --- | --- | --- |
| Donor # | Age | Sex | Race | Pain history | DRG level | Cause of death |
| 1 | 12 | F | white | none reported | L4 | anoxia |
| 2 | 47 | M | white | potential pain | L4 | ICH (stroke) |
| 3 | 5 | M | white | none reported | L4 | anoxia (seizure) |
| 4 | 53 | F | white | reported pain | L4 | ICH (stroke) |
| 5 | 53 | M | white | none reported | L4 | CVA (stroke) |
| 6 | 26 | M | white | reported pain | L4 | Head trauma (MVA) |
| 7 | 27 | F | unknown | none reported | L1 | Head trauma (MVA) |
| 8 | 34 | M | white | potential pain | L5 | Head trauma (MVA) |
| 9 | 25 | F | black | none reported | L4 | Head trauma (MVA) |
| 10 | 34 | F | hispanic | potential pain | L4 | anoxia (seizure) |
| 11 | 44 | F | middle-eastern | reported pain | L4 | CVA/SAH |
| 12 | 11 | M | white | none reported | L4 | head trauma (gunshot) |
| 13 | 54 | F | white | none reported | L4 | anoxia (OD) |
| Intact vs damaged grey matter in spinal cord |  |  |  |  |  |  |
| Donor # | age | sex | race | grey matter damage | DRG | Cause of death |
| 12 | 11 | M | white | N | L5 | head trauma (gunshot) |
| 14 | 28 | M |  | N | L4 |  |
| 15 | 49 | F |  | N | L4 |  |
| 16 | 20 | F |  | N | L5 |  |
| 17 | 28 | M | black | Y | L5 | anoxia (cardiac arrest) |
| 18 | 27 | F | unknown | Y | L4 | anoxia (drowning) |
| 19 | 46 | M | black | Y | L4 | anoxia (OD) |

During the course of these experiments, we also collected spinal cord from these donors and observed gray matter damage in spinal cord from a subset of patients (**Figure 1E**). This tissue damage correlated with extended time on life support prior to organ donation. We found that β*-NRXN* SS4 expression was significantly reduced in hDRG from donors with damaged spinal cord tissue compared to age- and sex-matched controls (**Figure 1F-G**). There was also a small, but not significant, reduction in SS4 expression in α-*NRXN* isoforms (**Figure 1F**), but we likely do not have sufficient statistical power from these limited sample numbers. Importantly, we were able to perform patch-clamp recordings on hDRG isolated from these donors for separate studies. This suggests that human DRG neurons were viable and healthy, and is consistent with our experience and the work of others that hDRG obtained from organ donors are particularly resilient (Davidson et al., 2014; Han et al., 2015; Payne et al., 2015; Xu et al., 2015; Zhang et al., 2015; Valtcheva et al., 2016; Cai et al., 2021; Tavares-Ferreira et al., 2022; Zurek et al., 2024). This initial observation of *NRXN* SS4 exclusion in human nervous system tissue from these donors prompted us to investigate if neurexin alternative splicing in DRG was indeed plastic using a reverse translational approach in rodent models.

### Activity-dependent changes in neurexin alternative splicing

We first asked if *Nrxn* alternative splicing at SS4 was regulated by activity in mouse DRG neurons. In contrast to most neurons in the CNS, DRG neurons do not form synaptic connections *in vitro* and are largely quiescent. We treated cultured DRG neurons with a depolarizing stimulus of 30 mM KCl or 1 µM PGE_2_ an inflammatory mediator that sensitizes DRG neurons (**Figure 2A**) (Davidson et al., 2014; McIlvried et al., 2025). DRG cell numbers and axon growth labeled by βIII-tubulin and NF200 were similar across both treatment conditions relative to controls (**Figure 2B**) and we have previously shown that treatment with 30 mM KCl does not cause cell death in DRG neurons(McIlvried et al., 2025). We designed similar primers for each mouse *Nrxn* isoform and used the same approach to quantify SS4 inclusion. There was a significant reduction in SS4 alternative splicing following KCl treatment for both alpha and beta isoforms of *Nrxn1* and *Nrxn3* relative to saline controls, while *Nrxn2* transcripts remained unchanged (**Figure 2C-I**). On the other hand, treatment with PGE_2_ did not alter *Nrxn* SS4 ratios compared to DMSO controls (**Figure 2C-I**). This result demonstrates that membrane depolarization reduces SS4 incorporation in DRG neurons, similar to the activity-dependent shift to SS4- *Nrxns* in cerebellar neurons (Iijima et al., 2011).

**Figure 2:**
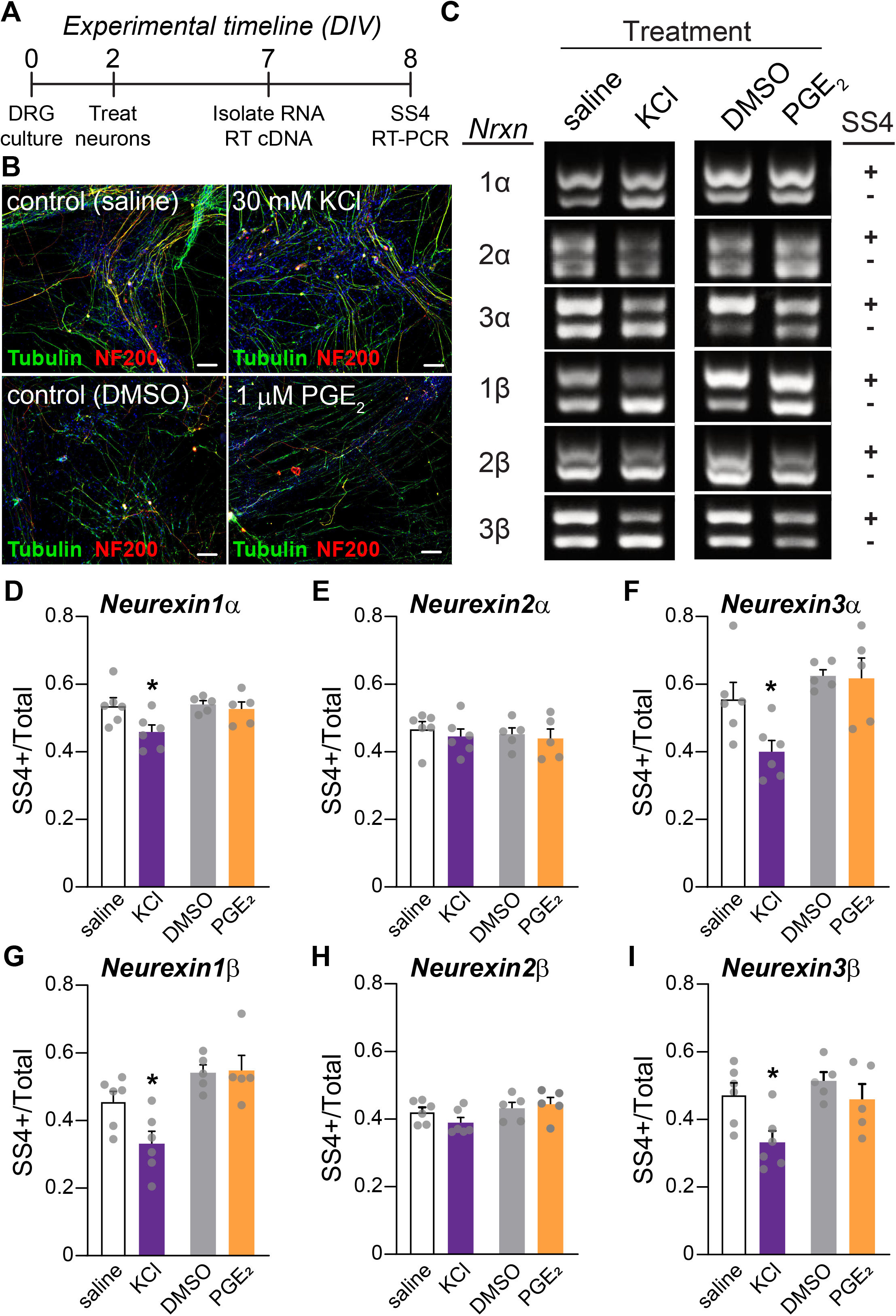
**Depolarization promotes *Nrxn* SS4 exclusion *in vitro.*** (A) Experimental timeline for *in vitro* experiments using cultured mouse DRG. DIV: days *in vitro* (B) Confocal images of cultured DRG neurons immunostained for the pan-neuronal marker βIII-tubulin (green), NF200 (red) for myelinated neurons and DAPI (blue). Neurons were treated with 30 mM KCl or saline control, and 1 µM PGE_2_ or DMSO control. Scale bar is 100 µm. (C) Representative images of ethidium bromide-stained gels from RT-PCR products of SS4+/- alternative splicing bands for the indicated *Nrxn* isoforms. (D-I) Summary graphs of the SS4+/Total quantification for the indicated alpha (D-F) and beta (G-I) neurexins. N=6 separate cultures for saline and KCl treatment, and N=5 for DMSO and PGE_2_. *p<0.05 using unpaired t-tests.

We next asked if *Nrxn* alternative splicing could be regulated by changes in DRG neuron activity *in vivo*. We chose the complete Freund’s adjuvant (CFA) model of inflammatory pain which produces an increase in spontaneous and evoked activity, primarily in C-fiber nociceptors (Djouhri et al., 2006; Weng et al., 2012). After baseline testing for mechanical sensitivity, we injected either CFA or saline (control) bilaterally into the hindpaws of adult mice (**Figure 3A**). We confirmed mechanical hypersensitivity at 24 hours that lasted for at least 7 days (**Figure 3B**). After behavioral testing, we purified RNA from L3-5 DRG and performed similar RT-PCR analysis. We did not find any changes in *Nrxn* alternative splicing at SS4 across any of the isoforms at either 24 hours or 7 days following CFA (**Figure 3C-H**), indicating that these splicing changes are not driven by acute inflammatory pain *in vivo*.

**Figure 3:**
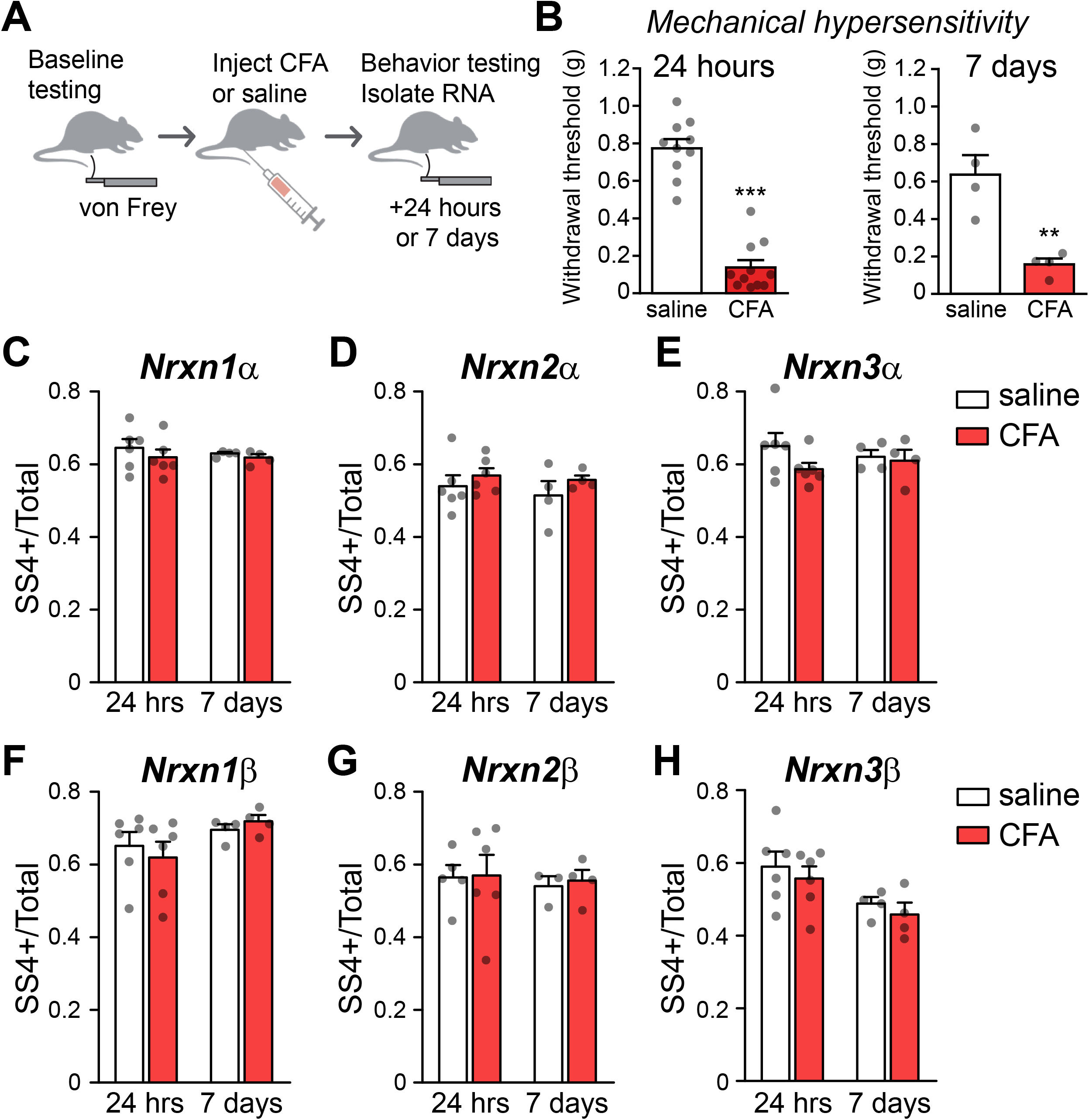
Inflammatory pain does not alter *Nrxn* SS4 alternative splicing. (A) Cartoon of the experimental design for testing *Nrxn* alternative splicing *in vivo* using the complete Freund’s adjuvant (CFA) model of inflammatory pain in the mouse hindpaw. (B) Summary graphs of persistent hindpaw mechanical hypersensitivity to von Frey filaments induced by CFA after 24 hours (left) and 7 days (right). (C-H) Summary graphs of the SS4+/Total quantification for the indicated alpha (C-E) and beta (F-H) neurexins 24 hours and 7 days after CFA injection. n=4-6 for saline and CFA-treated mice. **P<0.01, ***P<0.001 using unpaired t-tests.

### Neurexin alternative splicing following dorsal rhizotomy

In cultured DRG neurons, we observed a reduction in SS4 expression across all *Nrxn* isoforms that plateaued after 72 hours. We hypothesized that this could be due to the loss of synaptic connections, since cultured DRG neurons do not form synapses with each other *in vitro*, or an injury response to axotomy. To determine if either of these possibilities are relevant to *Nrxn* alternative splicing *in vivo*, we performed bilateral dorsal rhizotomies in adult mice to sever the central axon branches from L3-5 DRG (Basbaum, 1974) (**Figure 4A**). One week after dorsal rhizotomy, mice were ambulatory but began to develop lesions on the denervated hindpaws. We isolated RNA from the L3-5 DRG that underwent the rhizotomy or sham surgery, along with the L1,5,6 DRG to serve as an additional control, and performed RT-PCR to assess SS4 exon splicing (**Figure 4B**). We observed significant reductions in SS4 expression in L3-5 DRG for *Nrxn-1*α, *Nrxn-2*α, and *Nrxn-2*β but not in the other isoforms compared to sham operated mice (**Figure 4C-H**). We did not observe splicing changes for any *Nrxn* isoform between groups in the uninjured L1,5,6 DRG (data not shown). While this experiment is unable to separate whether these observed changes were due to decreased synaptic connectivity, nerve damage, or perhaps both, it does indicate that *Nrxn* alternative splicing is plastic *in vivo*.

**Figure 4:**
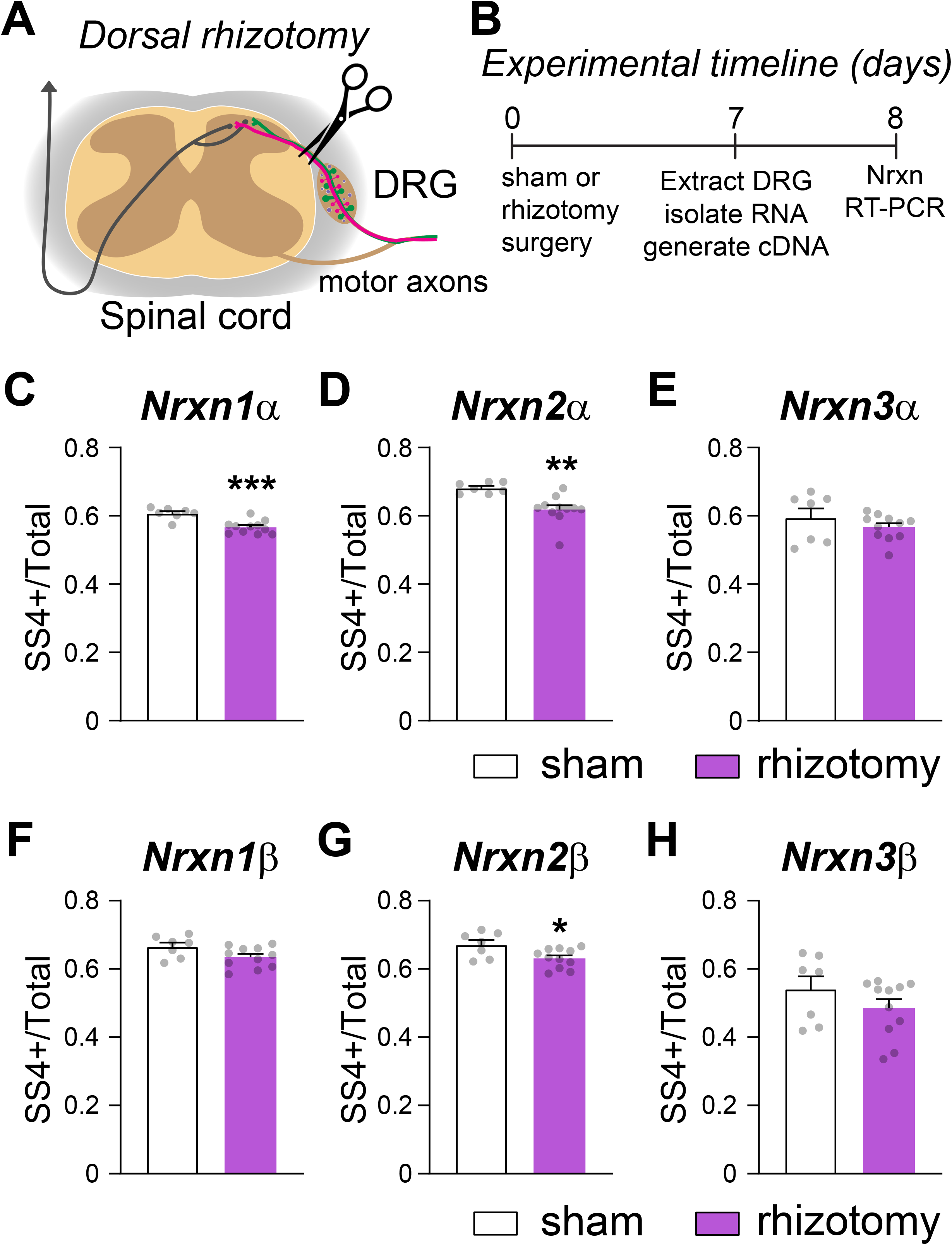
Dorsal rhizotomy promotes SS4 exclusion. (A) Cartoon of the dorsal rhizotomy approach to sever central axonal projections of DRG into the spinal cord. (B) Experimental timeline for testing *Nrxn* SS4 splicing after dorsal rhizotomy. (C-H) Summary graphs of the SS4+/Total quantification for the indicated alpha (C-E) and beta (F-H) neurexins one week after rhizotomy or sham surgeries. N=11 mice for rhizotomy and n=7 mice for sham surgery. *p<0.05, **p<0.01, ***p<0.001 using unpaired t-tests.

### Neurexin alternative splicing during somatosensory circuit development

Since *Nrxn* alternative splicing influences binding affinities to a variety of potential postsynaptic ligands (Südhof, 2017; Gomez et al., 2021), we asked if SS4 inclusion was plastic during somatosensory circuit development. Nearly all studies to date have focused on the role of neurexins in the brain, so we first compared *Nrxn* alternative splicing patterns in adult mouse DRG, spinal cord (SC) and cortex (Cx) to determine if there are tissue-specific differences in SS4 inclusion. We found that alternative splicing of *Nrxn* transcripts at SS4 were surprisingly consistent at approximately 60% SS4+ inclusion across all isoforms in adult mouse DRG (**Figure 5A-F**). *Nrxn* splicing at SS4 was similar between mouse DRG and SC for *Nrxn1* and *Nrxn2*, but there was a higher proportion of SS4+ *Nrxn3* transcripts in SC compared to DRG or cortex (**Figure 5A-F**). Overall SS4 inclusion was lower in cortex compared to DRG or SC across all *Nrxn* isoforms.

**Figure 5:**
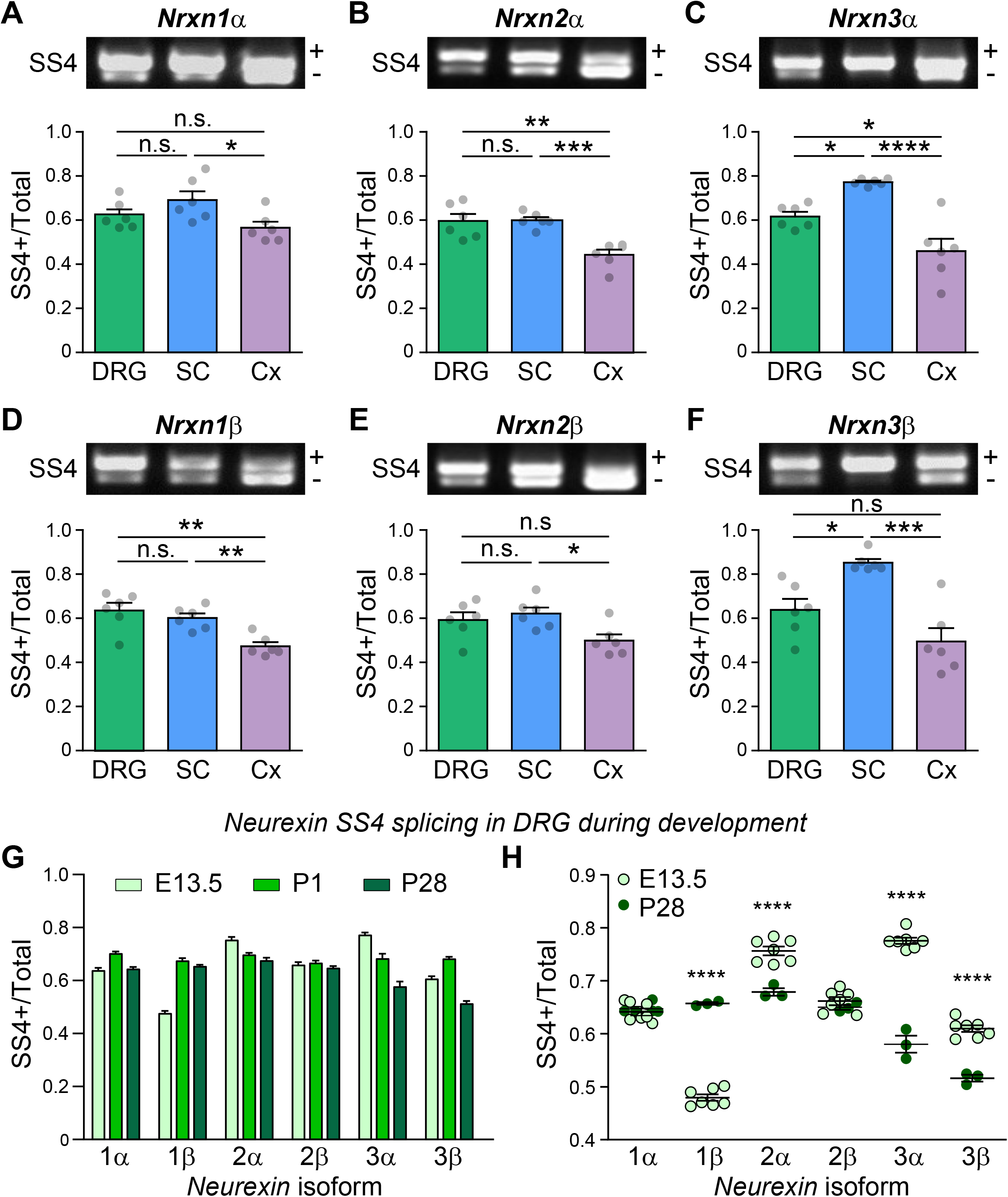
***Nrxn* alternative splicing at SS4 is dynamic in DRG neurons during somatosensory circuit development.** (A-F) Representative image of SS4+/- RT-PCR products and summary graphs of SS4+/Total expression for alpha- (A-C) and beta-neurexins (D-F) in dorsal root ganglia (DRG), spinal cord (SC) and cortex (Cx) from adult mice. N=6 mice for each tissue. (G) Summary graphs of SS4 inclusion across all neurexins in DRG from embryonic day 13.5 (E13.5), postnatal day 1 (P1), and postnatal day 28 (P28) mice. Statistical significance was tested using one-way ANOVAs and there is a significant difference across age groups for all *Nrxn* isoforms except *Nrxn2*β. Individual differences between ages using Tukey’s multiple comparisons test are omitted for clarity. (H) Comparison of SS4 inclusion for each *Nrxn* isoform between E13.5 and P28 mice. Statistical significance was tested using one-way ANOVAs with Tukey’s multiple comparisons across all three ages and significant differences are indicated between E13.5 vs P28. n=7 E13.5 mice, n=5 P1 mice, and n=3 P28 mice. n.s. not statistically significant, *p<0.05, **p<0.01, ***p<0.001, ****p<0.0001 for all graphs.

Lastly, we tested whether *Nrxn* alternative splicing at SS4 was plastic during somatosensory circuit development. We quantified SS4 inclusion ratios in mouse DRG at embryonic day 13.5 (E13.5) when DRG axons have extended to, but not yet innervated, the spinal cord (Mirnics and Koerber, 1995; Mirnics and Koerher, 1995; Hasegawa et al., 2007), postnatal day 1 (P1) when spinal cord innervation is complete, but synapses are still immature (Fitzgerald, 2005; Hasegawa et al., 2007) and P28 when these circuits are mostly mature (Meltzer et al., 2021). We found that SS4 exon inclusion in *Nrxn1*α and *Nrxn3*β exhibited a transient increase from E13.5 to P1 before returning to embryonic levels by P28 (**Figure 5G,H**). *Nrxn1*β SS4+ transcripts increased substantially between E13.5 to P1 and remained constant at P28 (**Figure 5G-H**). SS4 inclusion in *Nrxn2α* and *Nrxn3α* gradually decreased from E13.5 to P28, while alternative splicing at SS4 remained unchanged in *Nrxn2β* (**Figure 5G,H**). This result indicates that *Nrxn* alternative splicing is dynamic during somatosensory circuit wiring and may regulate synaptic connectivity and synapse maturation of the peripheral somatosensory system.

### Neurexin alternative splicing differs between species

Based on the splicing differences of *Nrxn* isoforms we observed across the nervous system in mouse, we last asked if *Nrxn* SS4 alternative splicing was different between human and mouse DRG neurons. We found that human DRG had a higher proportion of SS4 inclusion for longer α–Nrxn isoforms (**Figure 6A-C**) but observed the opposite result for the shorter β-Nrxns, particularly *NRXN3*β (**Figure 6D-F**). Interestingly, *NRXN1*α SS4 alternative splicing was the only isoform that did not exhibit significant species differences (**Figure 6A**). This is the only isoform that has been previously studied in native tissue between species and was found to have similar exon usage patterns between mouse and human neurons in the prefrontal cortex (Flaherty et al., 2019). These findings suggest that Nrxn alternative splicing and function may be more divergent across species than previously recognized.

**Figure 6:**
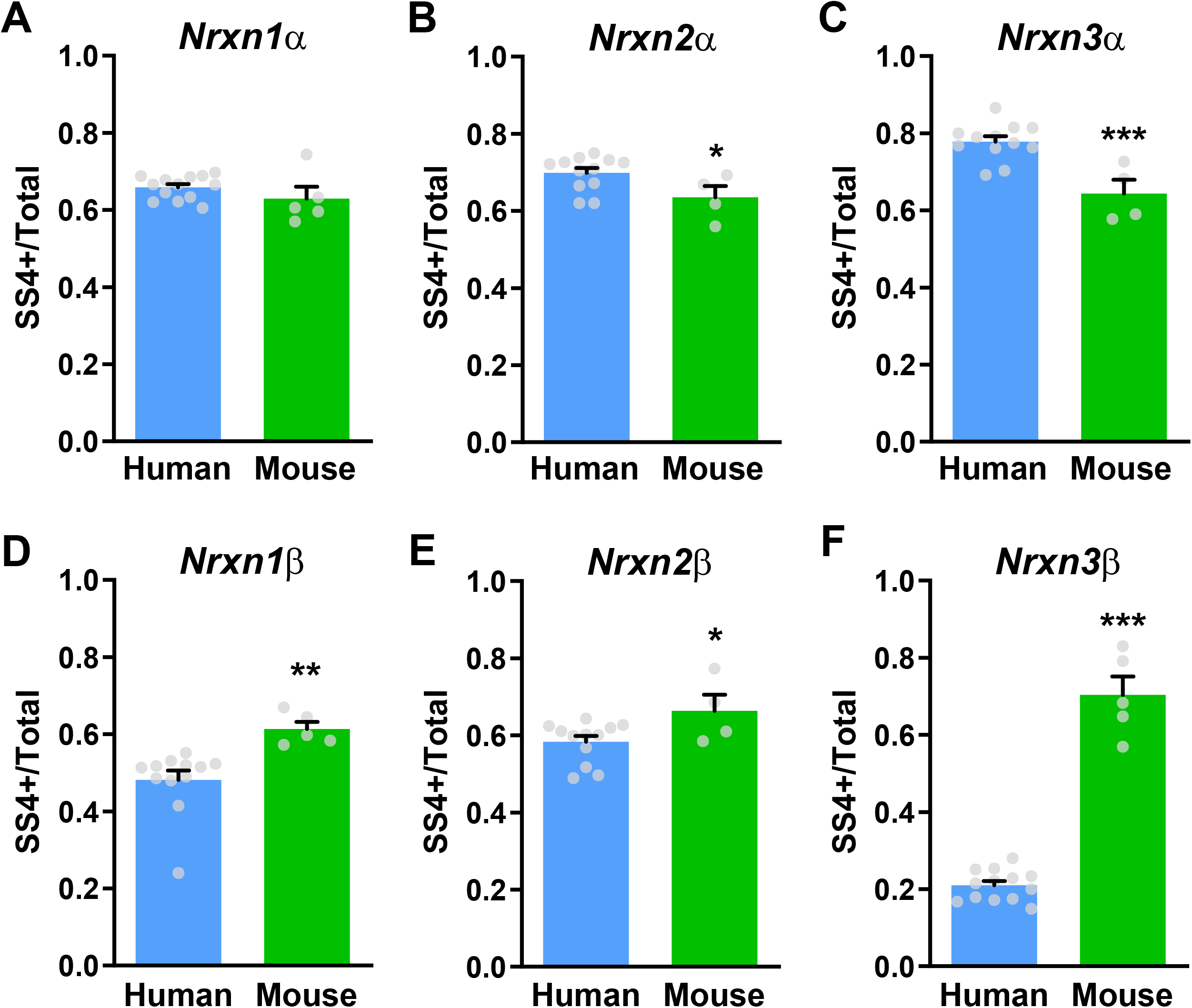
Divergence in *Nrxn* alternative splicing between mouse and human. Summary graphs of SS4+/Total expression for alpha (A-C) and beta-neurexins (D-E) between human and mouse DRG neurons. N= 12-13 (human) and 4-5 (mouse). Statistical significance was tested using unpaired t-tests, scholar*p<0.05, **p<0.01, ***p<0.001.

## Discussion

Here we show that *NRXN* alternative splicing at splice site 4 (SS4) is dynamic in both human and mouse somatosensory neurons. While we did not observe any changes in *NRXN* SS4 splicing from patients with prior pain history, we did observe a decrease in SS4+ transcripts in DRG from patients that had damaged spinal gray matter, suggesting that exon usage at this site may be plastic in human neurons. Taking a reverse translational approach using mouse models, we found that depolarization, but not sensitization with PGE_2_, decreased *Nrxn* SS4 usage in cultured DRG neurons. While we did not observe any changes in *Nrxn* alternative splicing at SS4 in the CFA model of inflammatory pain, we found that SS4 inclusion in *Nrxn1*α, *Nrxn2*α and *Nrxn2*β was reduced in DRG following rhizotomy of the central roots. *Nrxn* SS4 splicing patterns differed across mouse cortex, spinal cord and DRG, and was plastic in DRG during development. Lastly, we found that SS4 exon usage was conserved between mouse and human DRG neurons only for *NRXN1*α, with a higher proportion of SS4 inclusion in human *NRXN2*α and *NRXN3*α, but reduced incorporation of this exon for all human β- *NRXN* isoforms. These findings demonstrate that Nrxn splicing plasticity can be engaged by multiple mechanisms and may impart different synaptic properties between mouse and human at somatosensory synapses in the spinal cord.

Neurexins interact with a diverse array of potential postsynaptic ligands including neuroligins (Nlgn1-4), leucine-rich repeat transmembrane proteins (LRRTM1-4), latrophilin adhesion GPCRs, GABA_A_ receptors, and calsyntenins, in addition to secreted cerebellins (Cbln1-4), CA10/11, C1qls and neurexophilin proteins (Südhof, 2017; Gomez et al., 2021). The decrease in SS4+ transcripts we observed in response to depolarization and rhizotomy would be expected to shift binding away from cerebellins (Uemura et al., 2010; Elegheert et al., 2016) and towards dystroglycans (Sugita et al., 2001; Trotter et al., 2023), latrophilin adhesion GPCRs (Boucard et al., 2012), and the LRRTM family of postsynaptic cell adhesion molecules (Ko et al., 2009; Siddiqui et al., 2010). This picture is more complicated for the neuroligin family, where neurexin alternative splicing at SS4 modulates binding affinities, which can be additionally regulated by neuroligin alternative splicing (Boucard et al., 2005; Comoletti et al., 2006; Koehnke et al., 2010).

The molecular diversity generated by neurexin alternative splicing has been proposed to impart a code for specifying neuronal connections and synaptic properties (Südhof, 2017; Gomez et al., 2021). Different populations of DRG neurons encode distinct modalities of sensory information through cell-specific expression of ion channels and receptors and have distinct innervation patterns to relay this information to appropriate targets in the spinal cord. To understand the physiological implications of *Nrxn* alternative splicing, transgenic mouse lines now enable constitutive exon inclusion (Traunmüller et al., 2016a; Hauser et al., 2022; Lu et al., 2023), and cell-specific SS4 exclusion (Aoto et al., 2013; Dai et al., 2019). At CA1 to subiculum projections in the hippocampus, inclusion of SS4 in Nrxn1 selectively enhances NMDAR, but not AMPAR, synaptic currents (Dai et al., 2019). However, incorporation of SS4 in Nrxn3 suppresses AMPAR-mediated responses without affecting synaptic NMDARs, and prevents the induction of LTP (Aoto et al., 2013; Dai et al., 2019 p.2). In contrast, genetic deletion of the RNA-binding protein Slm2, which leads to predominantly SS4+ Nrxn isoforms, enhances AMPAR synaptic currents and prevents the induction of LTP at Schaffer collateral projections to CA1 (Traunmüller et al., 2016a). Thus, even in the same brain region, *Nrxn* alternative splicing exhibits isoform and cell-specific effects on synaptic function. Given the cell type-specific patterns of *Nrxns* and their alternative spliced variants in the brain(Fuccillo et al., 2015; Földy et al., 2016; Furlanis et al., 2019; Lukacsovich et al., 2019), it is interesting to speculate that similar cell type diversity of *Nrxn* expression might exist in DRG neurons. While this can be challenging to assess using relatively short reads from standard transcriptomic sequencing (Lukacsovich et al., 2019), recent advances in single-cell long-read sequencing could help answer these questions in both mouse and human sensory neurons (Arendt-Tranholm et al., 2024; Joglekar et al., 2024).

In this study we analyzed the net change in *Nrxn* SS4 alternative splicing from all DRG subtypes, which may have occluded our ability to resolve more dynamic changes in distinct cell types. In general, we observed an overall decrease in exon incorporation at SS4, but this varied across *Nrxn* isoforms and experimental manipulations. We do not know if particular DRG cell types are preferentially impacted in response to changes in activity, nerve damage, or during development. While we did not observe any changes in *Nrxn* alternative splicing in acute inflammatory pain, this largely drives increased activity only in C-fiber nociceptors innervating the hindpaw (Djouhri et al., 2006), which may have been minimally represented in our analysis of the entire ganglia. We also cannot say if the SS4 expression patterns we observed following rhizotomy are due to decreased synaptic connectivity, a response to nerve injury, or perhaps both. It will be interesting to test if different types of peripheral nerve injury, commonly used to model neuropathic pain in rodents, induce changes in *Nrxn* alternative splicing. Given that *Nrxn* SS4 expression regulates LTP in the hippocampus, this could potentially underlie some aspects of central sensitization at primary afferent synapses in the spinal cord in neuropathic pain. However, in contrast to hippocampal circuits, LTP at a subset of C-fiber synapses in the spinal cord can be induced by low frequency stimulation (LFS) around 2 Hz (Ikeda et al., 2006; Drdla-Schutting et al., 2012; Kim et al., 2015), which typically produces LTD at most synapses. In lamina I neurons, LTP requirements vary based on their projection targets in the brain (Ikeda et al., 2003, 2006), raising the possibility that differences in *Nrxn* alternative splicing might also regulate plasticity rules at specific synaptic connections in the spinal cord.

During development, we observed dynamic changes in *Nrxn* SS4 expression in DRG neurons. This is in contrast to developing mouse brain, where *Nrxn* gene expression and alternative splicing in distinct cell types is established by ∼E18 and stable into adulthood (Lukacsovich et al., 2019). DRG neurogenesis is complete by E13.5 (Ma et al., 1999; Landy et al., 2021) but neurons continue to undergo postnatal transcriptional maturation towards their final functional identity (Sharma et al., 2020). We observed splicing changes in *Nrxn* SS4 expression from P1 to adulthood, a period in which peripheral targets are still being innervated and responses to somatosensory stimuli are still being tuned. This plasticity in *Nrxn* alternative splicing may enable continued synaptic refinement as these circuits mature.

The changes in *Nrxn* alternative splicing we observed across different experiments raises the question of how this process is regulated in DRG neurons. Several RNA binding proteins including NOVA2, PTBP2, RBFOX1 and the STAR (signal transduction activators of RNA) family have been identified as regulators of *Nrxn* alternative splicing (Gehman et al., 2012; Wamsley et al., 2018; Saito et al., 2019). Three closely related STAR family members, SAM68, SLM1 and SLM2, all bind to *Nrxn* pre-mRNAs upstream of the SS4 site to generate *Nrxn* variants lacking this alternatively spliced exon (SS4-) (Iijima et al., 2011, 2014; Danilenko et al., 2016; Nguyen et al., 2016). These three proteins also exhibit cell-specific and mutually exclusive expression patterns in the hippocampus and striatum (Traunmuller et al., 2014; Gokce et al., 2016; Traunmüller et al., 2016a) and could function to generate specific *Nrxn* splice variants in different populations of somatosensory neurons.

The RNA-binding protein SAM68 is required for activity-dependent changes in *Nrxn* alternative splicing at SS4 in cerebellar neurons, and knockout mice have neurodevelopmental and motor deficits (Iijima et al., 2011). In the hippocampus, genetic deletion of SAM68 leads to exclusive generation of SS4+ *Nrxn* transcripts, which increases basal excitatory transmission and prevents the induction of NMDAR-dependent LTP. These mice also exhibit altered binding interactions between presynaptic neurexins and a variety postsynaptic adhesion molecules (Traunmüller et al., 2016b). The synaptic and behavioral deficits in SAM68 knockouts were rescued in *Nrxn1* transgenic mice that normalize SS4 splicing ratios. This suggests that SAM68 may regulate the activity-dependent and/or developmental changes in *Nrxn* alternative splicing in DRG neurons that we observed here. Because *Nrxn* alternative splicing influences binding affinity to potential postsynaptic adhesion molecules, it will be interesting to investigate somatosensory circuit development and nociceptive plasticity in these knockout animals.

We initially set out to test whether *NRXN* SS4 expression was altered in human DRG from donors with reported pain histories. While we did not find any differences across donors regardless of age, sex, or pain history, this study was spurred by the changes we found in human DRG from donors with damaged spinal gray matter. We do not know if the reductions in *NRXN* SS4 expression from these donors is a physiological response to decreased connectivity with spinal neurons or the result of neuronal cell death. However, our initial finding in human DRG revealed that *Nrxn* alternative splicing is plastic in somatosensory neurons and demonstrates the utility of a reverse translational approach in model organisms to gain deeper insights into observations of the human nervous system.

## Methods

### Human tissue collection

Human tissue was obtained from postmortem organ donors in collaboration with Mid-America Transplant as described previously (Valtcheva et al., 2016). DRG and spinal cord were transferred on ice in carbogenated modified aCSF (Ting et al., 2014) containing (in mM): 93 N-methyl-D-glucamine, 2.5 KCl, 1.25 NaH2PO4, 30 NaHCO3, 20 HEPES, 25 glucose, 5 ascorbic acid, 2 thiourea, 3 Na pyruvate, 10 MgSO4, 0.5 CaCl2, and 12 N-acetylcysteine; adjusted to pH 7.3 with HCl, 300-310 mOsm) and transported back to the lab within 2 hours after aortic cross-clamp. Tissue was then dissected and either used for culture or placed in RNALater (Thermo) for at least two days, prior to long-term storage at -80°C. Consent of tissue donation for research and medical information was obtained from next-of-kin by Mid-America Transplant. Donor demographics and medical history are provided in **Table 1**.

### Animals and procedures

All experiments in mice were approved by the Washington University School of Medicine Animal Care and Use Committee and in accordance with the National Institute of Health guidelines. Wild-type male and female C57Bl/6 mice were used for all experiments. Embryonic tissue was collected from prenatal mice generated from timed matings, with vaginal plug formation defined as embryonic day E0.5. Postnatal day P5-10 mice were used for neuron cultures. 8–12-week-old mice were used for all surgical procedures and behavioral testing. All mice were group-housed with *ad libitum* access to food and water and maintained on a 12 h light:dark cycle.

### RNA isolation and RT-PCR

RNA was isolated using TRIzol reagent (Themo-Fisher 15596018) according to manufacturer protocols. Cultured cells were rinsed once with PBS and lysed directly in TRIzol. Tissue was homogenized in TRIzol using dounce glass homogenizers until evenly disrupted. After a 5-minute incubation, 1/5 volume chloroform was added to the samples for another 3 minutes at room temperature. Samples were centrifuged at 12000 xg for 15 minutes at 4C. Supernatants were then transferred to new tubes and 0.5 ml of isopropanol (per 1 ml of TRIzol) was added, followed by 10-minute incubation at 4°C. Samples were centrifuged at 12000 xg for 15 minutes at 4°C and supernatants discarded. Pellets were then washed twice in 80% ethanol, centrifuged at 7600 xg for 4 min at 4°C before being resuspended in nuclease-free water. RNA yield and purity was quantified using a Nanodrop and either used immediately or stored at -80°C. cDNA was made from RNA using a reverse transcription kit (Thermo-Fisher 4368814) according to manufacturer instructions using equal amounts of RNA for all samples.

RT-PCR was performed using 2 ul of each cDNA input and Taq polymerase (Promega). All primers sequences (IDT) recognize both mouse and human *NRXN* mRNA except *NRXN3*α, in which human and mouse specific forward primers were designed. Primer sequences are (all 5’ to 3’): *Nrxn1*α forward (ATGGGATGGCTTCAGCTGTG), *Nrxn1*β forward (GGTCACCAGCATCCTTGC), *Nrxn1* reverse (CCCGCCAATTATTATGGTTGC), Nrxn2α forward (ACTTCCTATGGAGGCCCTGT), *Nrxn2*β forward (CCACCACGTCCACCACTT), *Nrxn2* reverse (TGGCTGTTGAAGATGGTCAG), Human *NRXN3*α forward (ATGATGCTCTTCATCGGAGC), mouse *Nrxn3*α forward (ACGATGCTCTCCACAGGAGT), *Nrxn3*β forward (TCTACACCTGGCCAGCAAAT), *Nrxn*3 reverse (AGAGAGTTGGCCTTGGAAGA), *Gapdh* forward (CCCTTCATTGACCTCAACTACATG), *Gapdh* reverse (TGGTGAAGACGCCAGTAGACTC). 10 ul of each RT-PCR reaction was separated on a 2% agarose gel, stained with ethidium bromide, and imaged. All PCR products matched their expected size (SS4- and SS4+, in bp): *Nrxn1*α (466 and 556), *Nrxn1*β (434 and 524), *Nrxn2*α (413 and 503), *Nrxn2*β (468 and 558), *Nrxn3*α (632 and 722), *Nrxn3*β (398 and 488).

### Primary DRG culture

DRG cultures were prepared from postnatal day 0-10 mice. DRG were dissected in HBSS with 10 mM HEPES (HBSS+H). Tissue was enzymatically digested with papain (45U, Worthington, cat. # LS003126) for 20 min at 37°C, rinsed in HBSS+H, followed by collagenase (1.5 mg/mL; Sigma, cat. #C6885) for 20 min at 37°C. Ganglia were washed and resuspended in culture media consisting of Neurobasal A (GIBCO), 5% FBS (Life Technologies), 1x B27 (GIBCO), 2 mM glutamax (Life Technologies) and 100 µg/mL penicillin/streptomycin (Life Technologies). Neurons were dissociated by mechanical trituration through glass pipettes, passed through 40 µm filters, and plated at a density of 40,000 cells/well onto poly-D-lysine and collagen coated coverslips.

### Immunocytochemistry and confocal imaging

Culture DRG neurons were fixed in 4% PFA on ice for 10 minutes and washed 3x with PBS. Cells were permeabilized and blocked for 1 hour with 0.3% TritonX-100, 3% goat serum in PBS followed by incubation in a 1:1000 dilution of primary antibodies (rabbit βIII- tubulin, Covance, cat. no. MRB-435P, RRID: AB_663339; mouse NF200, Millipore cat. no. MAB5266, RRID: AB_2149763) overnight at 4°C. After 3 washes in PBS, coverslips were incubated in 1:2000 dilution of secondary antibodies (donkey anti-rabbit AF488 and goat anti-mouse AF568, both from Life Technologies) for 1 hour at room temperature. Coverslips were mounted with Prolong Gold (Life Technologies) and imaged on a Leica SPE confocal microscope.

### Rhizotomy

Adult C56Bl/6 mice (8-12 weeks old) were anesthetized with 3% isoflurane and after shaving the fur and sterilizing the site, an incision was made over the lumbar spinal column. After performing a laminectomy, the dorsal roots from L3-5 DRG were carefully cut bilaterally using spring scissors ∼5 mm from the dorsal root entry zone. The incision was closed using metal surgical staples and mice were transferred to a heating pad to recover from anesthesia. Sham surgeries consisted of the full laminectomy, but dorsal roots were left intact.

### Behavior testing

Mice were briefly anesthetized with isoflurane and 10 µl of either saline or CFA (1 mg/ml; Sigma) was injected bilaterally into the plantar surface of the hindpaw. After acclimation, mechanical sensitivity was tested using the up-down method with von Frey filaments. The 50% withdraw frequency was calculated from the average of 3 trials. Baseline measurements were taken prior to CFA injection, and the development of mechanical hypersensitivity was tested 24 hours and 7 days after injection.

### Experimental design and statistical analysis

Experimenters were blinded to the treatment groups during testing and analysis. The ratios of *Nrxn* SS4 alternative splicing were quantified using Fiji software (NIH). Images were background subtracted and the SS4/Total ratio was calculated by dividing the integrated density for the larger SS4+ PCR product by the summed integrated density from both SS4+ and SS4- products. Data was analyzed in Excel and graphs and statistical analysis were generated using GraphPad Prism software. Unpaired two-tailed t-tests were used for comparing two groups, while one-way ANOVAs with Tukey’s multiple comparisons tests were used for comparing more than two groups. Statistical significance was determined by P<0.05. All graphs represent the mean ± standard error of the mean, with individual data points plotted.

## Acknowledgements

We sincerely thank the donors and their families for their generous gifts to science that made this research possible. We thank the staff at Mid-America Transplant for providing access to donor tissue and their facilities, especially Dr. Gary Marklin and Nicole Mullins for assistance with donor coordination, and John Lemen for technical support with tissue extraction. We also thank Dr. Robert Gereau for suggestions and comments on this manuscript and Sherri Vogt for technical assistance. This work was supported by NIH R01 NS130046 (BAC), the American Pain Society Future Leaders in Pain Research Award (BAC), the McDonnell Center for Cellular and Molecular Neurobiology (BAC and RWG) and the NIH PRECISION Human Pain Network U19 NS130607 (BAC and RWG).

## Author contributions

JJY designed and performed research, analyzed data, made figures, and edited the manuscript. BAC designed research, analyzed data, made figures, and wrote the manuscript.

## Conflict of interest

The authors declare no competing financial interests.

## Notes

### Competing Interest Statement

The authors have declared no competing interest.

